# Imaging Mass Cytometry (IMC) as a Tool to Characterize Circulating Tumor Cells (CTCs) in Preclinical Mouse Models

**DOI:** 10.64898/2025.12.18.695262

**Authors:** Milind Pore, Kuppusamy Balamurugan, Abigail Atkinson, Devynn Breen, Paul Mallory, Ashley Cardamone, Lois McKennett, Christine Newkirk, Shikha Sharan, William Bocik, Esta Sterneck

**Affiliations:** Imaging Mass Cytometry Laboratory, Frederick National Laboratory for Cancer Research, Leidos Biomedical Research, Inc., Frederick, MD, USA; Cancer Innovation Laboratory, Center for Cancer Research, National Cancer Institute, Frederick, MD, USA; Laboratory Animal Sciences Program, Frederick National Laboratory for Cancer Research, Leidos Biomedical Research, Inc., National Cancer Institute, Frederick, MD, USA

**Keywords:** imaging mass cytometry, imaging-proteomics, circulating tumor cells, liquid biopsy, xenograft

## Abstract

Circulating tumor cells (CTCs), particularly multicellular clusters, are associated with poor prognosis and may provide insight into mechanisms of metastasis and therapy resistance. Unbiased approaches for functionally characterizing CTCs in liquid biopsies are therefore urgently needed. Here, we evaluate multiplex imaging mass cytometry (IMC) for CTC analysis in mice bearing human xenograft tumors. In a single-step workflow, IMC uses metal-conjugated antibodies to simultaneously detect numerous proteins and post-translational modifications in minimally processed, small-volume blood samples collected from the tail vein or heart. Using breast cancer cell lines and a patient-derived xenograft (PDX), we assessed a panel of antibodies, including human-specific markers such as Lamin B1 (LMNB1), to enable cross-species interpretation. Combined with manual review, HALO AI–based cell segmentation was used to identify CTCs and quantify marker expression. This approach enables studies of how genetic and pharmacologic interventions alter the properties of single CTCs and CTC clusters in tumor-bearing mice.

## Introduction

Metastasis is in part due to circulating tumor cells (CTCs), especially CTC clusters, which are associated with poor prognosis and treatment resistance [1-3]. CTCs increase in numbers after biopsy, surgery, and during neo-adjuvant treatment [4], leading to their exploration as therapeutic targets[5]. CTCs can be assessed with liquid biopsies, which hold immense promise for cancer patients by enabling non-invasive, real-time monitoring of tumor dynamics [6]. While many advancements in techniques to isolate and/or enumerate CTCs have been made, investigations of biological mechanisms are very challenging due to their scarcity. Almost all CTC detection methods rely on two steps, first to enrich the CTCs from the millions of blood cells and then specifically isolate/detect the CTCs based on protein marker expression. In addition, the enrichment and isolation techniques that are required for the study of CTCs in clinical samples invariably expose the cells to physical manipulations and stresses which may alter characteristics of the cell [7, 8]. Because of the high sensitivity of nucleic acid-based methodologies, most characterizations have been conducted at the level of genomics and transcriptomics compared to proteomics [9], although proteins are the most functionally relevant entities. Detection of proteins is especially relevant for insights into protein modifications or subcellular localization. Lastly, while CTCs from patients may provide valuable correlative information, only preclinical mouse models permit genetic and pharmacological manipulations that will lead to mechanistic insights and facilitate the development of novel therapeutics [10].

Fluorescent immunostaining has been used for limited targeted investigations of CTCs, but large-scale proteomic approaches remain lacking [30, 31]. In addition, the number of markers that can be analyzed simultaneously is limited with this technology. Multiplex applications require repeated cycles of staining and washing, which risk losing these rare single cells during analysis. In contrast, multiplex imaging mass cytometry (IMC) of metal-labeled antibodies allows for the simultaneous detection of more than 40 protein epitopes in situ in an exclusive single-step process [11]. Though IMC is mostly employed for spatial analysis of single cells in tissues [12, 13], a small number of studies have applied the technology to CTCs from patients [14, 15]. Here, we evaluated 31 metal-labeled antibodies for detection and characterization of specifically human breast cancer CTCs in mouse blood without prior enrichment of CTCs. Antibodies were assessed with cell lines and cell line-spiked mouse blood. Xenograft mouse models were used to detect and characterize the CTCs, including single-cell marker quantification, in part through AI-assisted technologies. Limitations of the technologies and needs for future developments are being discussed. The panel presented here can be expanded for specific research questions and as an unbiased tool to assess CTC protein biology. Developing this tool for the functional characterization of single CTCs versus CTC clusters in pre-clinical models will shed light on the biology of this elusive but critical cancer cell population, and lead toward a better assessment of the efficacy and clinical utility of therapeutic drugs [16].

## Materials and Methods

### Cell lines and culture

MCF-7 cells were obtained from ATCC; SUM149 and SUM159 cells originated from Asterand Bioscience (New York, USA). IBC-3 cells were kindly provided by Dr. Junichi Kurebayashi (Kawasaki Medical School) through Dr. Wendy A. Woodward (MDACC) and MDA-MB-231-LM2-GFP-Luc cells (termed (MB-231-LM2) by Dr. Joan Massague. Cell lines were authenticated in 2022 and 2025 by GenePrint®10 (Promega, Wisconsin, USA), and *Mycoplasma* tested by qPCR. Cells were cultured in a 5% CO_2_ incubator at 37°C in media with antibiotics as follows: MCF-7, MDA-MB-231-LM2, and MDA-MB-468 cells were in Dulbecco’s Modified Eagle’s Medium, MCF-7 in addition with 1 mM sodium pyruvate; SUM159 in RPMI with 2 mM glutamine, 10 mM HEPES, 1 mM sodium pyruvate, 1X nonessential amino acids (GIBCO, #11140-050) and 55 mM β-mercaptoethanol (GIBCO, #21985-023); SUM149, and IBC-3 in Ham’s F-12 media (GIBCO, #31765092) with 1 μg/ml hydrocortisone and 5 μg/ml Insulin; Fetal bovine serum (FBS) was added at 10% except for SUM159 (5%). Cell culture grade chemicals were from Sigma Aldrich unless indicated otherwise. As indicated, 5×10^4^ cells were seeded on Millicell EZ Slides (Millipore, cat number PEZGS0816) and analyzed after 24-48 h.

### Western blotting analysis

Whole cell extracts were prepared and Western blotting performed as described [14] with primary antibodies from Cell Signaling Technology (Pan-keratin #4545T; EpCAM #93790T; Vimentin #5741S), and Santa Cruz Biotechnology (GAPDH #sc32233).

### Xenograft mouse models

The breast cancer PDX BCM-5471 originated from the PDX Core of the Baylor College of Medicine [17] and was propagated in NOD/SCID/Il2rg−/− (NSG) mice essentially as described [17]. PDX fragments or 2×10^6^ cells of the indicated cell lines were implanted into the inguinal fat pads of 9–17-week-old female mice. Tumor volumes were calculated as V = (W(2) × L)/2. When tumors reached up to 2000 mm^3^, about 100 μl blood was collected from the tail vein and/or by cardiac puncture.

NCI-Frederick is accredited by the Association for Assessment and Accreditation of Laboratory Animal Care International (AALACi) and follows the Public Health Service Policy for the Care and Use of Laboratory. Animal care was provided in accordance with the procedures outlined in the Guide for the Care and Use of Laboratory Animals (National Academies Press, 2011) including those pertaining to studies of neoplasia (National Research Council, 1996). All experiments were conducted under protocols approved by the IACUC at NCI-Frederick.

### Blood collection and preparation for imaging analysis

Peripheral blood was collected from the tail vein into BD Microtainer tubes with K_2_EDTA (BD, REF 365974). For spiking, about 10^4^ tumor cells in 10 ul were added to 100 ul blood before erythrocytes lysis with 2 ml ACK lysis buffer (Lonza, #BP10-548E) followed by centrifugation at 1500 rpm for 5 min at 4°C. Cells were washed once with 2 ml of ACK lysis buffer and twice with PBS. For IMC analysis, cells were suspended in 50 μl PBS by gentle pipetting 2-3 times using 20-200 microliter tips, applied to 3-well adhesion slides (Superior Marienfeld #0900100 & 0906100) and allowed to be attached on glass surface for 1h at 37°C in a dry oven.

### Antibody-metal labeling and immunocytochemistry

Antibodies compatible with immunocytochemistry that were BSA-free and at 100 µg quantity with at least 0.5 mg/ml concentration were purchased from different vendors (Table S1). Metal labeling of antibodies was performed using Maxpar® X8 Multimetal Labeling Kit-40 Rxn (201300) as described by the manufacturer’s protocol (Standard BioTools). Cells on EZ slides were fixed with 2% paraformaldehyde for 20 min, followed by permeabilization with 0.1% Triton X for 10 min. Slides were blocked with 3% BSA in PBS for 1 h and stained with a mixture of metal-labeled antibodies at different dilutions (Table S1) overnight at 4^0^C. Slides were washed with PBS and counterstained with Cell-ID™ Ir191/193 genomic DNA intercalator (Standard BioTools, #201192B) at 1:500 dilution. Cells were washed with PBS followed by quick dip in ddH2O to remove the salts. Slides were dried and ablated with the Hyperion imaging system.

### Imaging mass cytometry

Data were acquired by laser ablation with a Hyperion XTi Imaging System (Standard BioTools). The instrument was tuned with 3-element tuning slides provided by the manufacturer prior to sample acquisitions. A minimum dual count of Lu 800, transient crosstalk <25% and resolution (first and second mass) >400 criteria were used to successfully pass the tuning of an instrument. For stages one and two, at least three ROIs of 1500*1500 µm size were ablated. For stage three, with detection of CTCs from xenograft and PDX models, the panorama was generated of smeared cells and the ROIs of 1500*1500 µm size were created to cover all the cells from the 3-well slides. The data obtained in .mcd format was uploaded on MCD viewer software (Standard BioTools) to check the staining and manually identify the CTCs. The CTCs were defined as DNA-IR+, CD45- and Pan-keratin+ cells. Pilot experiments had been performed on two earlier versions of Hyperion imaging system (Hyperion and Hyperion+) with similar results.

### Nuclear segmentation and automated CTC counting

StarDist, Cellpose, and HALO were used for nuclear segmentation. In QuPath, segmentation was performed with the pre-trained DSB2018 model in StarDist, which uses star-convex object detection [18, 19]. Additionally, segmentation using the pre-trained cyto3 model in Cellpose, a generalist segmentation algorithm, was also assessed in QuPath [20, 21]. The third method utilized the HALO Image Analysis platform (Indica Labs), which uses convolutional neural networks for cell level segmentation. Single cell quantification and CTC phenotyping was performed using the HALO HighPlex FL module v4.2.14 with the CTC phenotype defined to identify DNA3^+^Pan-keratin^+^CD45^-^ cells.

## Results

To assess the performance of specific antibodies for the evaluation of protein expression in human CTCs, we utilized three experimental approaches (**Figure 1)**. Stage 1 determined the antibody signals in cell lines cultured directly on slides. In stage 2, cell lines grown in cell culture dishes were spiked into mouse blood to assess staining performance under blood matrix conditions. These samples were spotted onto 3-well adhesion slides. Finally, blood from xenograft models was collected via tail vein and/or cardiac puncture for CTC detection and characterization.

**Figure 1.**
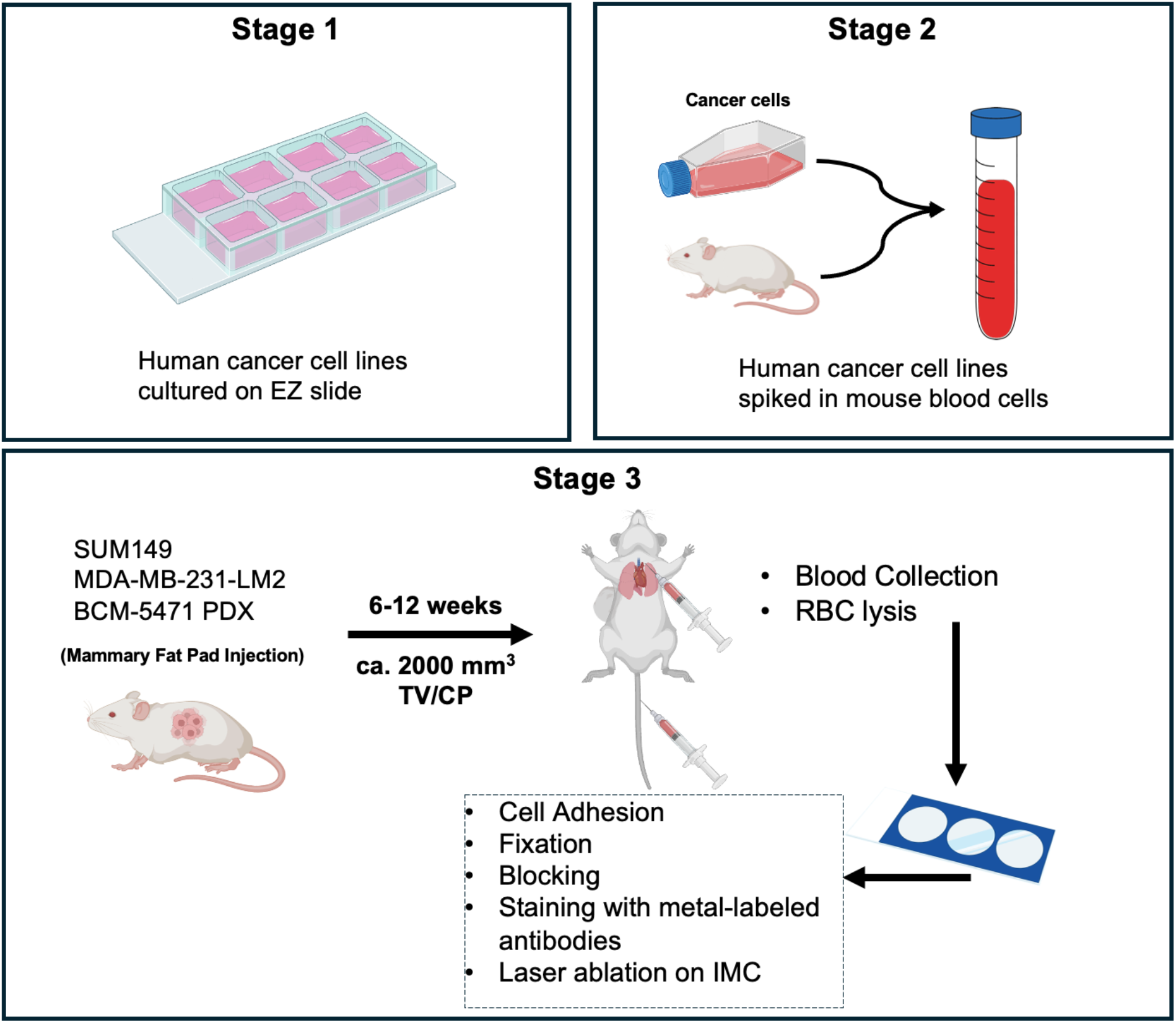
Schematic illustration of the experimental steps toward IMC analysis of CTCs. (Figure created in part in BioRender.com).

To select suitable cell lines, we determined the expression levels of the epithelial molecule EpCAM and the mesenchymal protein vimentin across a panel of breast cancer cell lines by Western blots (**Figure 2A**). In addition, we analyzed the reactivity of a Pan-keratin antibody, which produced most signal in epithelial cell lines and weak but detectable signals in mesenchymal tumor cells, as had been reported [22]. One mesenchymal (MB-231-LM2) and two epithelial (IBC-3, SUM149) cell lines were chosen to culture directly on glass chambered slides. Three days later, the slides were processed for metal-labeled antibody staining followed by laser ablation on the Hyperion imaging system. EpCAM and vimentin clearly distinguished the three cell lines. IBC-3 and MB-231-LM2 cells showed distinct subcellular abundance and distribution of Pan-keratin with SUM149 presenting a mixed phenotype (**Figure 2B**). These data suggested that the Pan-keratin antibody may be useful to identify most of these cells irrespective of epithelial/mesenchymal phenotype. We proceeded with SUM149 and MB-231-LM2 cells grown on Millicell EZ slides to assess a panel of metal-labeled antibodies (**Figures 3** and **Supplemental Figures S1-S2**, left column). Next, the cells were spiked into mouse blood before spotting onto adhesion slides (**Figures 3** and **Supplemental Figures S1-S2**, right column). We had pre-determined that plating 100 µl of nucleated cells from mouse blood in 15×15 mm wells provided sufficient spacing without cell crowding. Mouse blood without spiking was used as negative control and to determine cross-/reaction of antibodies with mouse cells. As shown before, SUM149 and MB-231-LM2 were distinguished by EpCAM and vimentin respectively. Most other markers showed some degree of variability of expression within the spiked cells in each cell line. Interestingly, both panels showed Pan-keratin negative cells (Figure 3, white circles), indicating that this is not a universal marker. Overall, signals were higher in stage 1 compared to stage 2, which could reflect biological differences due to different growth substrates or be due to the processing of cells for spiking. Nine antibodies showed satisfactory signals (Figure 3) at stage 1 and/or 2 that appeared specific based on negativity of mouse cells or selectivity for one of the two cell lines (Pan-keratin, EpCAM, Vimentin, EGFR, NOS2, CK8, SMAD2, CD44, ZEB1, plus CD45). Six antibodies generated high background, at least under the chosen conditions (**Supplemental Figure S1**) and 8 antibodies produced no or inconclusive signals which may also be due to genuine lack of expression (**Supplemental Figure S2 and Table S1**). Only one of two tested antibodies against E-cadherin (Table S1) gave reliable results in distinguishing the epithelial IBC-3 cells and the mesenchymal MB-231-LM2 cells (**Supplemental Figure S3**). Several other tested antibodies did not produce satisfactory signals in our hands including one against specifically human Na/K-ATPase α1 protein (Table S1). [23]

**Figure 2.**
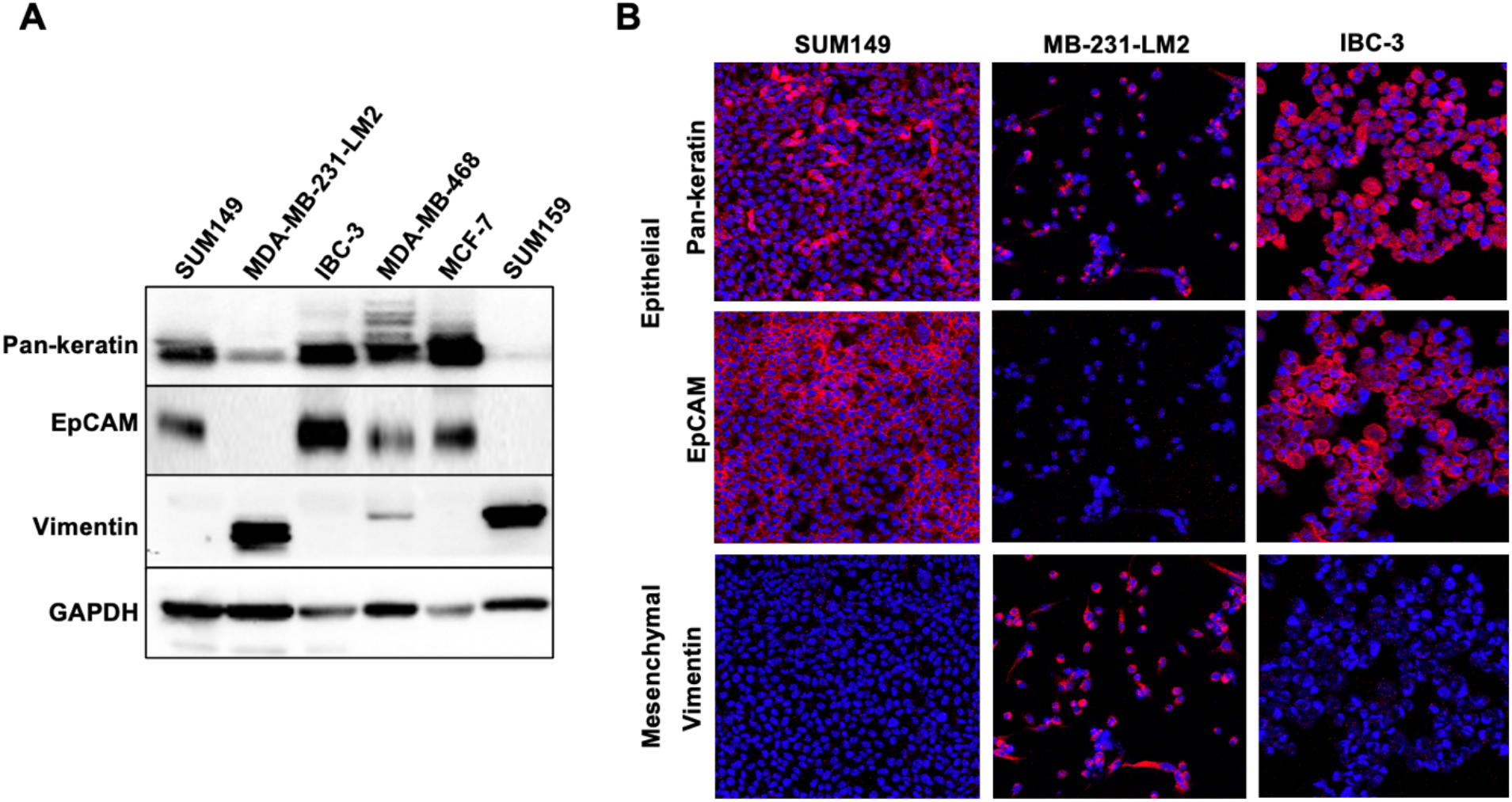
Comparison of Western and IMC analysis of specific antibodies. **(A)** Western blotting analysis of the indicated proteins in whole cell extracts of a panel of breast cancer cell lines. **(B)** Imaging mass cytometry images of proteins in a subset of cell lines grown on EZ slides. The same primary antibodies were used as in panel A, though labelled with unique metals showing red pseudo color (nuclei are shown in blue).

**Figure 3.**
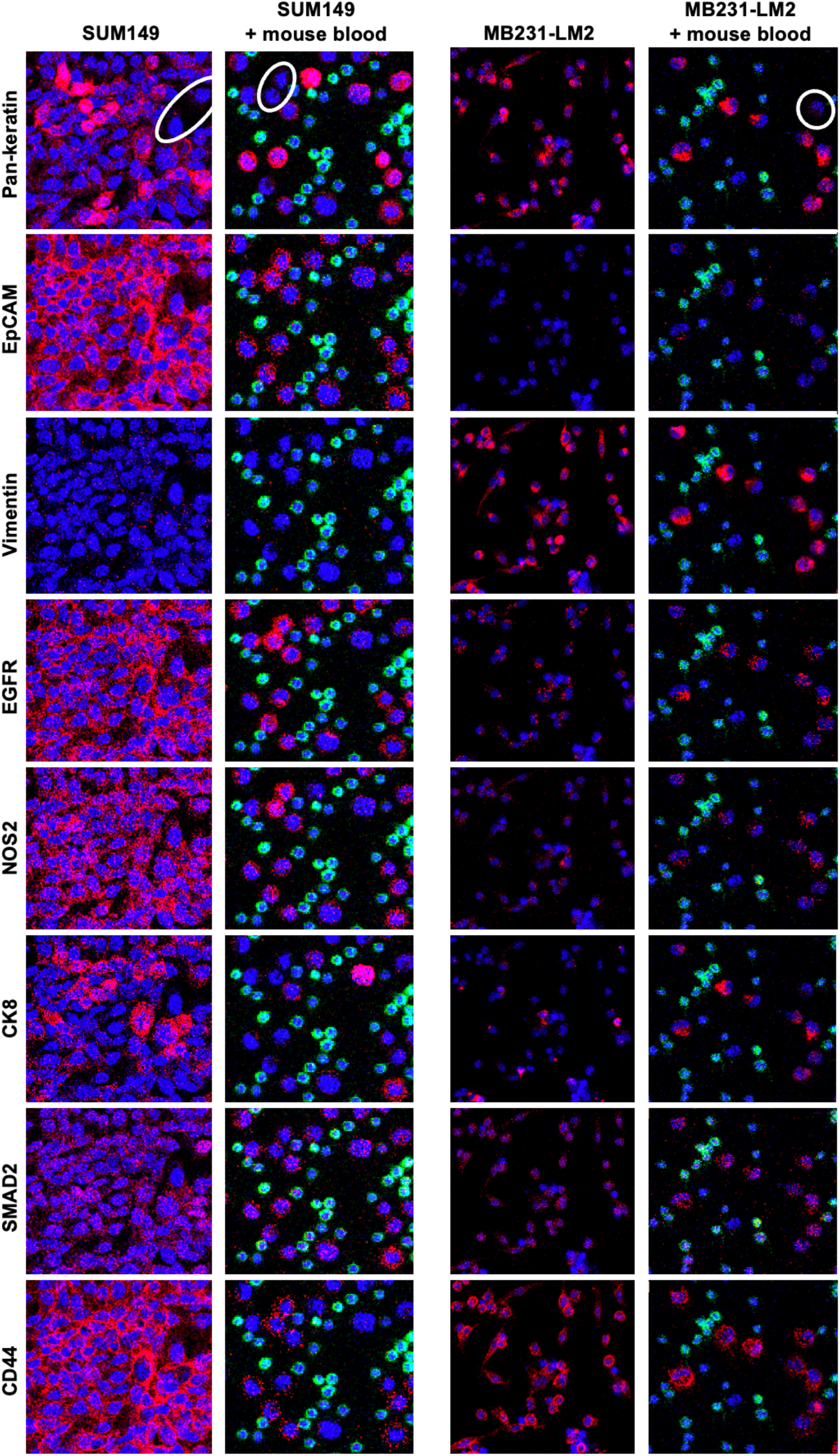
Comparison of IMC signals by specific antibodies in cell lines with and without admixed mouse blood. SUM149 (Left) and MB-231-LM2 (right) cells were spotted directly onto slides or first spiked into mouse blood as indicated. Nuclei are shown in blue and murine immune cells are identified in green by anti-CD45 staining.

In Stage 3, we sought to analyze CTCs in blood from mice with tumors derived from the SUM149 and MB-231-LM2 cell lines or the patient-derived xenograft BCM-5471. CTCs were identified manually by presence of a nucleus, Pan-keratin positivity and CD45 negative staining. Table S2 summarizes the frequency of CTCs in blood collected by tail vein and/or cardiac puncture. We observed a trend toward more CTCs in blood collected from the heart, which was however not statistically significant. **Figure 4** shows IMC data for three CTCs from each of the three cancer models. Some antibodies which had generated higher reactivity in stages 1-2, e.g. P-cadherin and CK18, showed more variable staining in CTCs. Conversely, some markers such as CK5 and CK19, were seen only in Stage 3 CTCs at varying intensities but generated no or weak signals with cultured cells (stage 1 and 2) suggesting that their expression may be induced in vivo. Despite the small number of examples, the data also showed significant variability in expression between CTCs in the same model, suggesting that weak signals may be interpreted as genuine. For example, SUM149 CTC#2 expressed ZEB1 and higher levels of CK5 than CTC1 and CTC3. In BCM-5471, only CTC#2 showed high levels of CD44 while CTC#3 was negative for most markers. In BCM-5471, SMAD2 was absent, nuclear, and cytoplasmic, respectively, in the three examples of CTCs. Comparison of two 2-3 cell clusters from BCM-5471showed similarities and contrasts (**Figure 5**). For example, EpCAM and vimentin showed more signal in Cluster#1 respectively Cluster #2, and CK19 was detected in only one cell of each cluster.

**Figure 4.**
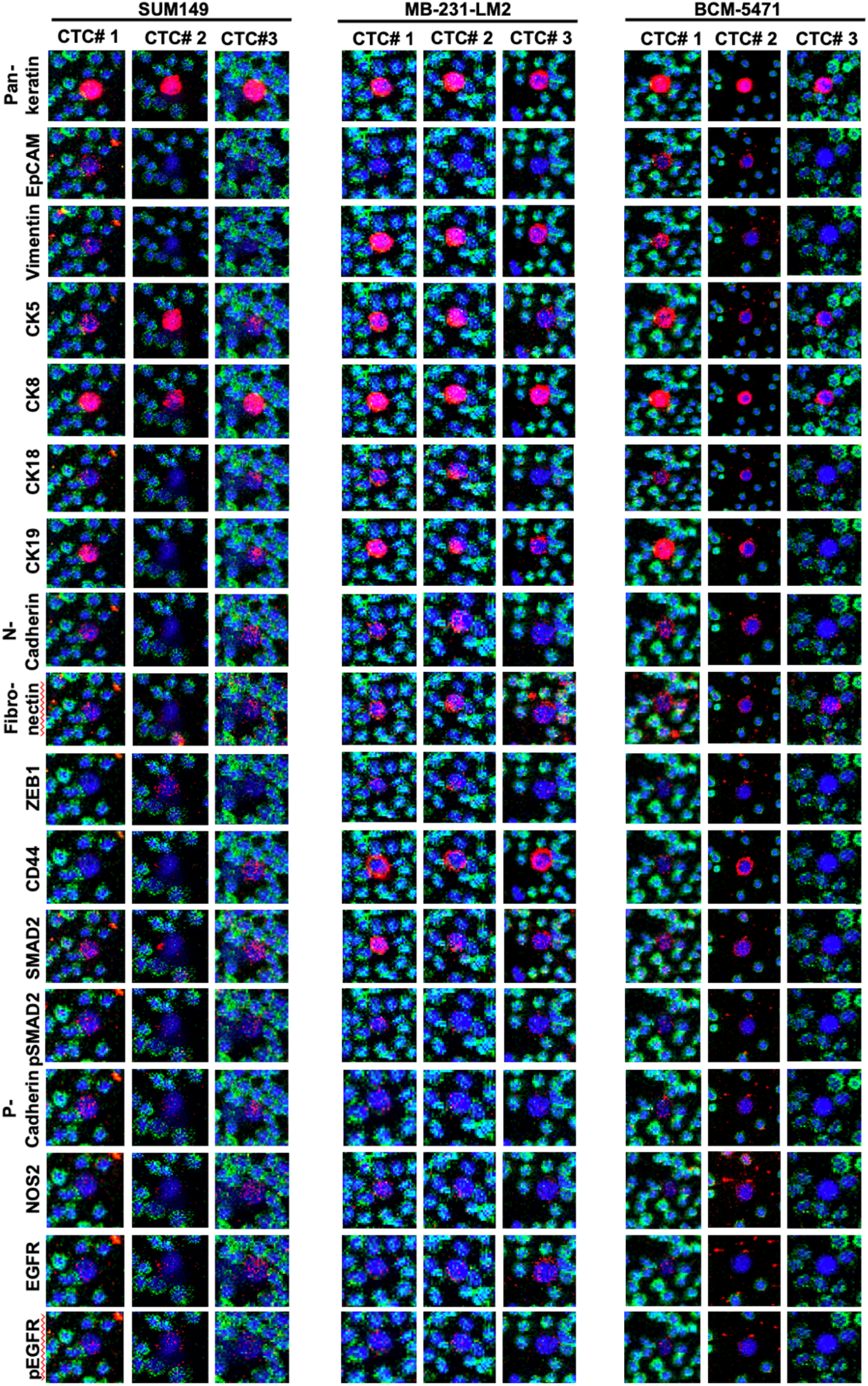
Comparison of IMC results across CTCs and mouse models. IMC images of antibodies against the indicated proteins in three different CTCs each from mice with SUM149, MB-231-LM2 or PDX xenograft tumors.

**Figure 5.**
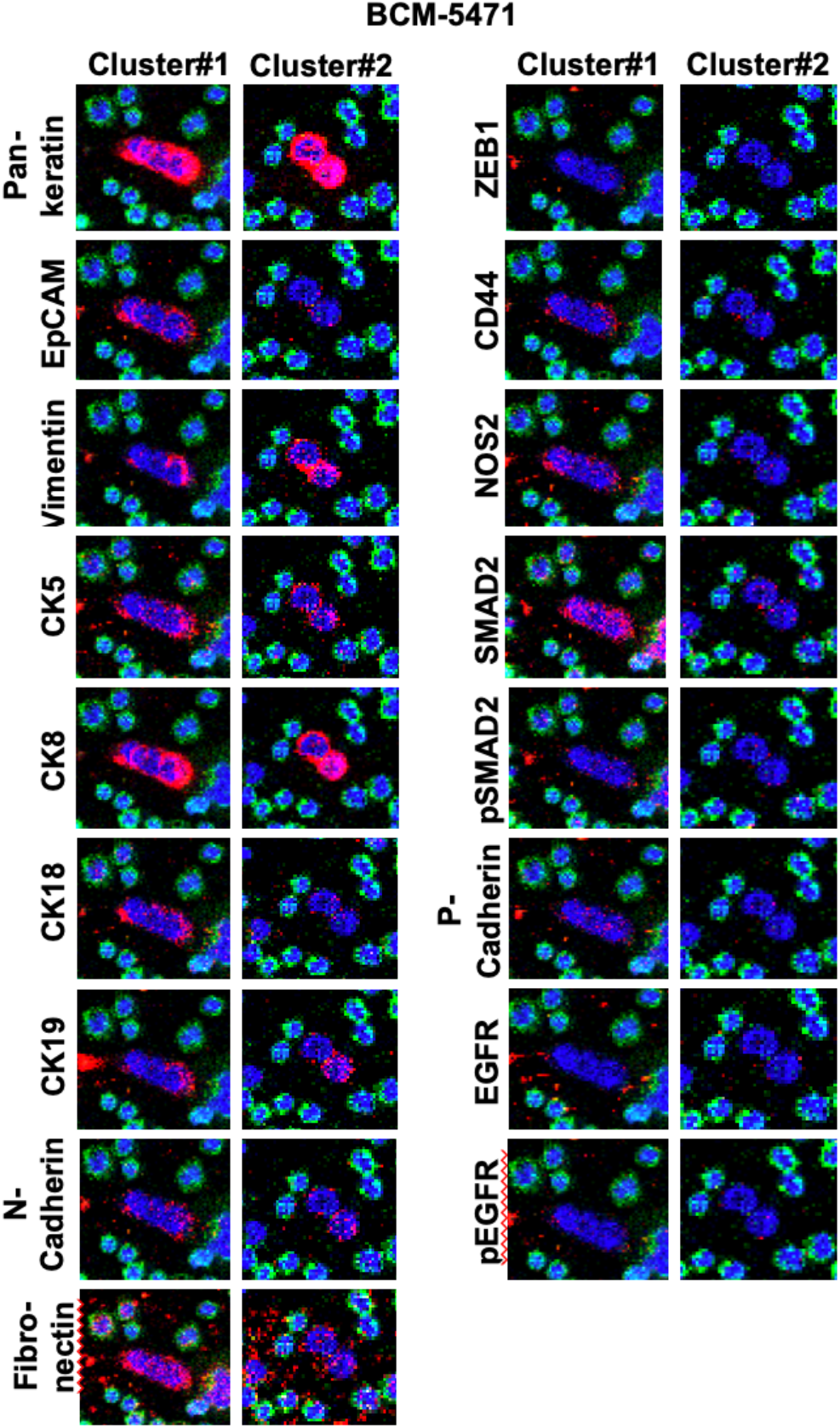
IMC analysis of CTC clusters from BCM-5471 PDX. IMC images of the indicated antibodies in two 2-3 cell clusters of CTCs from mice with BCM-5471 PDX tumors.

To move beyond labor-intensive manual enumeration and evaluation of CTCs, we also developed an automated cell segmentation and CTC detection pipeline. The results from three nuclear segmentation methods (HALO, Cell pose, and StarDist) were compared across multiple images for the three cancer cell models in the study (**Figure S4A**). In evaluating the accurately identified nuclei from the images surveyed, the HALO AI segmentation performed better than the StarDist and Cellpose pre-trained methods. The StarDist model had difficulty identifying nuclei in dense clusters while Cellpose missed detection of lower intensity nuclei. Moving forward with the HALO AI segmentation, over 2000 nuclei from the IMC images were annotated giving the algorithm substantial ground truth examples to train from. The training of the classifier was validated at every ten thousand annotations to evaluate the peak in its performance. At 40k iterations, the validation of several images showed the most accurate detection and segmentation of the nuclei, and therefore, this trained algorithm was chosen as the segmentation module to construct the analysis module. The acquisition of xenografts and PDX derived slides were ablated on different versions of Hyperion Imaging system including, Hyperion, Hyperion+ and the newest Hyperion XTi. This presented differences in the intensity of the nuclear marker, DNA3, which created challenges in using one consistent segmentation module for all images acquired. To accurately detect nuclei on images with lower signal intensities in this channel, HALO AI default pre-trained segmentation was utilized on several images in the analysis that the HALO custom AI underperformed on.

CTCs were identified by the presence of DNA3 and Pan-keratin, and the absence of the immune cell marker CD45 (**Figure S4B**). The HALO classification of the CTC phenotype was optimized by performing a visual inspection for Pan-keratin positivity. Due to the signal intensity differences between image sets and instruments, a general survey of cells expressing Pan-keratin was performed first. This preliminary analysis involved broad thresholds with a phenotype defined to include DNA3^+^/Pan-keratin^+^/CD45^-^ cells. The Pan-keratin signal intensity needed to be greater than 1.4 and more than 20 percent completeness, a parameter that requires a set percentage of the cell to be above the minimum positivity threshold, while the CD45 signal needed to be less than 2.0 and 20 percent completeness for the cell to be counted as a CTC. After a review of these preliminary results, thresholds for the DNA3, Pan-keratin, and CD45 were defined for the CTC phenotype per cell model and instrument as outlined in Table S3. Examples of the number of CTC counts with HALO AI cell segmentation classifier in comparison to manual CTC counts is shown in Table S4 and suggested overall concordance. Importantly, HALO AI segmentation enables the quantification of expression differences at the single cell level. To demonstrate this utility, **Figure 6** shows examples of quantifications of several signals in single and cluster CTCs from two mice with BCM-5471 tumors with one cluster CTC each. The data again illustrate significant variations between CTCs and can be used to generate hypotheses such as preferential co-expression of vimentin and cytokeratin 8 in cluster CTCs, suggesting a hybrid E/M phenotype which is highly relevant for metastasis [24].

**Figure 6.**
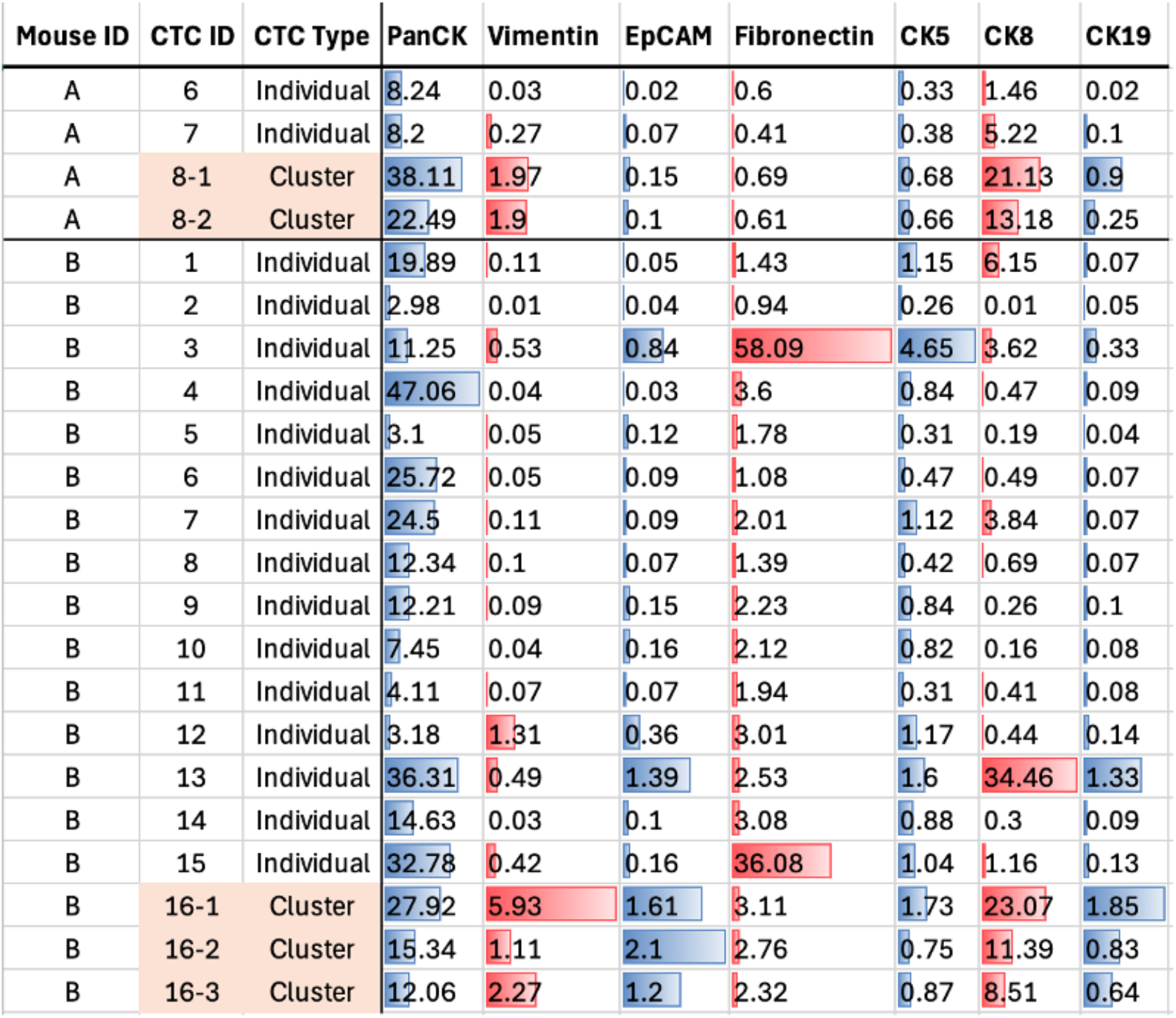
Quantification of IMC signals in single CTCs and CTC clusters. HALO-derived quantification of specific antibody signals for individual cells (CTC ID**#**) in CTC clusters and as single CTCs in blood of BCM-5471 tumor-bearing mice (values are mean marker intensities in arbitrary units).

Lastly, we sought to identify a universal human-specific marker because Pan-keratin is not reliable in detecting all cancer cells (see also Figure 3) and may also mark mouse epithelial or endothelial cells. Furthermore, expression of CD45 on CTCs has been described and, albeit rare, CD45-negativity among criteria to call CTCs would miss such “dual-positive” cells [23]. First, we tested antibodies against MHC-II and the mitochondrial protein NUMA which is suitable for the detection of human cells in tissue sections of mice [25, 26] (**Figures 7A-B**). In a mix of MB-231-LM2 and IBC-3 cells, anti-NUMA1 showed unspecific staining and high background in mouse blood cells. Anti-MHC-II did not stain any CD45-positive mouse cells and marked all CD45-negatiev cells, but with a high dynamic range. MHC-II has been described as both positive and negative prognostic marker and can be downregulated in the process of immune evasion [27-29] which may further hamper its utility for identification of CTCs. Next, we evaluated the nuclear protein Lamin B1. This protein was readily detected in all Pan-keratin positive mesenchymal MDA-MB-231 and epithelial IBC-3 cells when spiked into mouse blood but was not seen in CD45+ cells (**Figure 7C**). In SUM149 xenograft bearing mouse blood, anti-Lamin B1 marked Pan-keratin positive cells with different combinations of other markers, e.g. CK5 and CK18 respectively (**Figure 7D**). We conclude that Lamin B1 is a useful addition to the panel for the identification and characterization of human CTCs in mouse blood.

**Figure 7.**
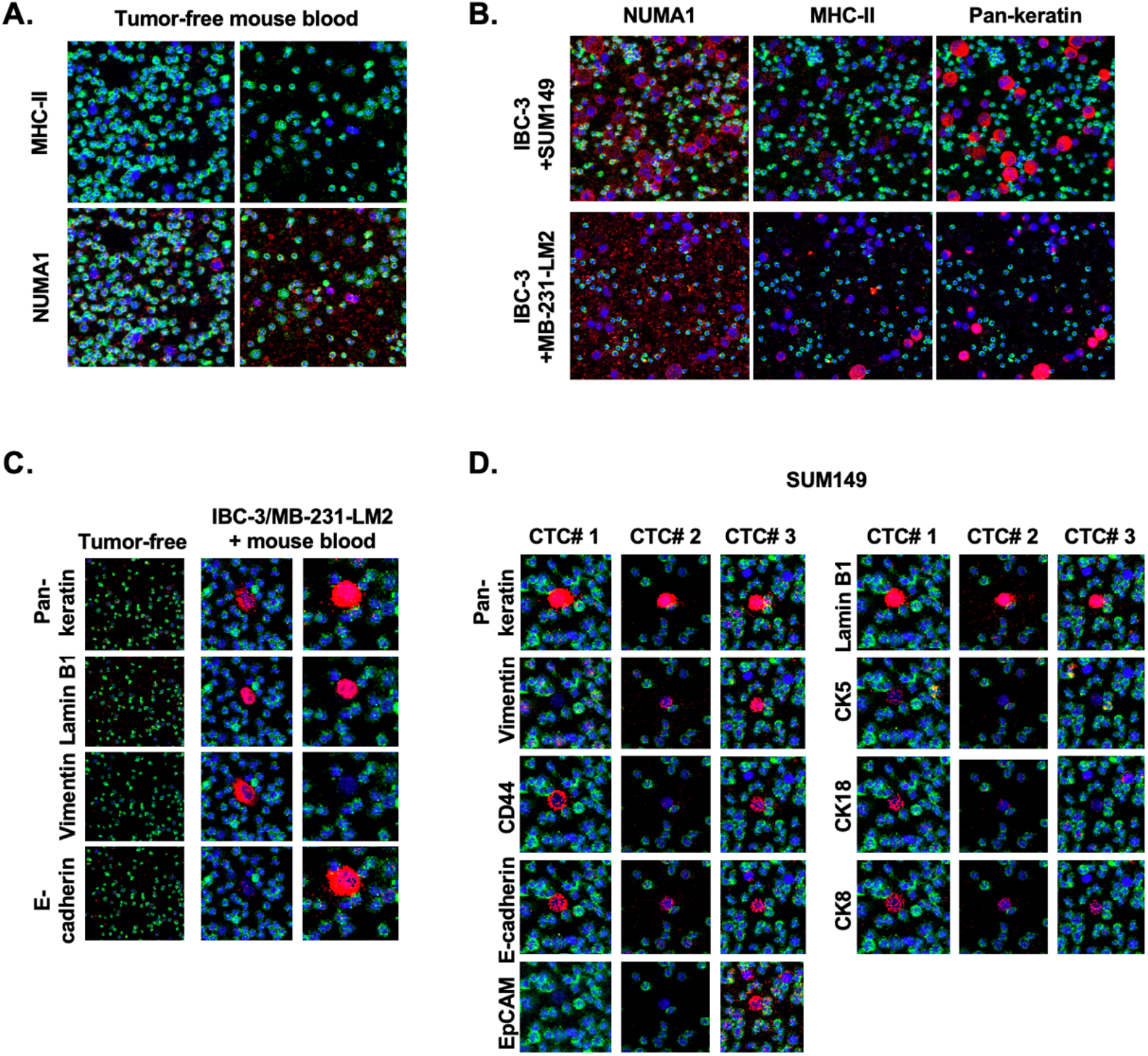
Evaluation of human-specific antibodies against MHC-II, NUMA1, and Lamin B1. IMC images of antibodies against the indicated proteins applied to (A) tumor-free mouse blood alone; (B,C) mouse blood alone and spiked with human cell lines as indicated; (D) three CTCs in blood from a SUM149 tumor bearing mouse.

## Discussion

Currently, enumeration is the only CTC application with a clear level of evidence for clinical use in liquid biopsy. However, in a survey of experts, phenotypic characterization of CTCs and biomarker expression for response prediction were considered the most important future clinical applications [16]. IMC has been explored for CTC characterization in patients but has not yet achieved wider application [14, 30]. Preclinical models therefore hold substantial promise for accelerating discovery of translationally meaningful CTC biology through mechanistic studies in mice. Here, we assessed IMC of CTCs in liquid biopsies from mice and provide a basis for further development of this methodology that may accelerate discovery and broaden the utility of CTCs in liquid biopsy research and application.

The key advantages of IMC lie in the use of over 40 metal-tagged antibodies with unique, non-overlapping spectra, and a single-step workflow. However, IMC also has limitations. Compared with multiplex fluorescence, it requires more expensive instrumentation and lacks signal amplification by secondary antibodies. The approach we took in stages 1-2, i.e. the validation of antibodies on cell lines proved useful in some but not all instances. A wider variety of cell lines, silencing of specific targets, or experimental treatments may be required to ascertain the specific of antibody signals. Antibodies that work in IMC on tissues may not work on liquid biopsies because of the differences in maple processing.

Furthermore, we found that CTCs can induce expression of proteins not detected in the same cells when cultured ex vivo. This possibility must be considered when validating antibody reactivity on cells that were grown ex vivo. In our hands, the mesenchymal MDA-MB-231-LM2 cells still reacted with anti-Pan-keratin. The subcellular distribution was droplet-like, as previously described for rhabdomyosarcoma cells [31] and a hepatocyte clone [32]. Therefore, the % cell coverage had to be adjusted for this cell line for the automated analysis workflow. For automated CTC detection, we recommend beginning with low thresholds followed by manual verification of called CTCs. Issues encountered at this step included Pan-keratin positivity “without nucleus” and “hybrid cells” positive for both CTC and immune-cell markers. The latter have been proposed to result from phagocytosis of human material by immune cells, cell fusion, exchange of proteins through cell-cell contact, or valid dual-positive CTCs [23] [33-35]. Because such cells were rare, their interpretation will depend on their relevance to the specific research question. Like many CTC studies [36], we initially relied on Pan-keratin to identify cancer cells. However, some cells were Pan-keratin-negative, suggesting that purely mesenchymal CTCs could be missed and that vimentin could be included in the initial CTC calling strategy. The use of Lamin B1 could mitigate this issue.

A general challenge in CTC research is distinguishing tumor cells from normal non-immune cells. In xenograft mouse models, human-specific antibodies against ubiquitously expressed proteins could help distinguish normal mouse-derived cells from human tumor-derived cancer cells. However, most antibodies against proteins of interest cross-react with both species. Of the three antibodies tested in this study, only Lamin B1 generated satisfactory specific signals in CTCs from a human cell line and a PDX model. Lamin B1 is abundantly expressed in breast cancers but is downregulated during senescence [37]. Although this limits its utility, senescent CTCs may not be relevant as metastasis-initiating cells. If this is a concern, other antibodies against ubiquitously expressed human proteins will need to be evaluated. Alternatively, genetically introduced protein tags or reporter genes could be used to confer cancer-cell specificity in xenograft mouse models.

Previously, it had been suggested that CTCs were undetectable in blood drawn from the tail vein, and the less feasible method of cardiac puncture was recommended for serial determination [38]. Our findings, together with those of a prior study [25], demonstrate that tail vein sampling is a viable method for monitoring CTCs and is therefore suitable for longitudinal studies. We also established that 100 µl of mouse blood is a sufficient biopsy volume for CTC detection across the tested models. Although CTC enrichment may be desirable, each isolation method has specific strengths and weaknesses, most require expensive equipment, and all involve cell manipulations that may alter cell biology [39]. We propose that cell spotting with minimal processing best preserves the original cellular phenotype. Future optimization could include comparing different slide-preparation approaches for CTC retention and evaluating quantitative retention relative to alternative detection methods.

In conclusion, we show that IMC can be a suitable single-step methodology for detecting and phenotypically characterizing CTCs in preclinical mouse models. The workflow presented here provides a platform for interrogating CTC heterogeneity, cluster formation, and dynamic biomarker expression. Applying this approach to CTC analysis in mouse liquid biopsies can support monitoring of genetic alterations and therapeutic interventions and help advance liquid biopsy applications beyond enumeration toward treatment-response prediction and precision oncology.

## Acknowledgements

This research was supported by the Intramural Research Program of the National Institutes of Health (NIH), ZIA BC 010307, and in part with federal funds under contract no. 75N91019D00024 including support from the Imaging Mass Cytometry Laboratory of the Frederick National Laboratory for Cancer Research. The contributions of the NIH author(s) were made as part of their official duties as NIH federal employees, are in compliance with agency policy requirements, and are considered Works of the United States Government. However, the findings and conclusions presented in this paper are those of the author(s) and do not necessarily reflect the views of the NIH or the U.S. Department of Health and Human Services.

AI tools were employed to refine English language usage during manuscript preparation.

## See Supplementary Material File for

### Supplementary Figures

**Figure S1: Comparison of IMC signals by specific antibodies in SUM149 cells with and without admixed mouse blood**. Representative images showing each target in red pseudo color, DNA in blue and CD45 in green.

**Figure S2: Comparison of IMC signals by specific antibodies in MDA-MB-231-LM2 cells with and without admixed mouse blood**. Representative images showing each target in red pseudo color, DNA in blue and CD45 in green.

**Figure S3: Validation of E-cadherin antibody**.

Expression of PanCK and E-cadherin as detected by clone 24E10 in MDA-MB-231-LM2 and IBC-3 cells individually (CD45 signal was omitted) and spiked together into mouse blood (CD45+ staining in green).

**Figure S4. Identification of CTCs by an automated and custom AI Halo cell segmentation classifiers. A)** Nuclear segmentation of liquid biopsies from three tumor cell models across three segmentation algorithms, HALO AI, Cellpose, and StarDist (scale bars, 50µm). White circles indicate regions with errors. **B)** Example of HALO segmentation images derived from the raw data as indicated (DNA: white, PanKeratin: red, CD45: green).

### Supplementary Tables

**Table S1**. Antibodies used or tested for IMC in this study.

**Table S2**. Comparison of the frequency of CTCs in blood collected by tail vein versus cardiac puncture.

**Table S3:** Threshold parameters for calling Pan-keratin+ CTCs by HALO HighPlex FL module per tumor model and instrument.

**Table S4:** Comparison of manual CTC counts versus identification by AI Halo cell segmentation classifier.

