## Supplementary Figures and Tables for "Imaging Mass Cytometry (IMC) as a Tool to Characterize Circulating Tumor Cells (CTCs) in Preclinical Mouse Models"

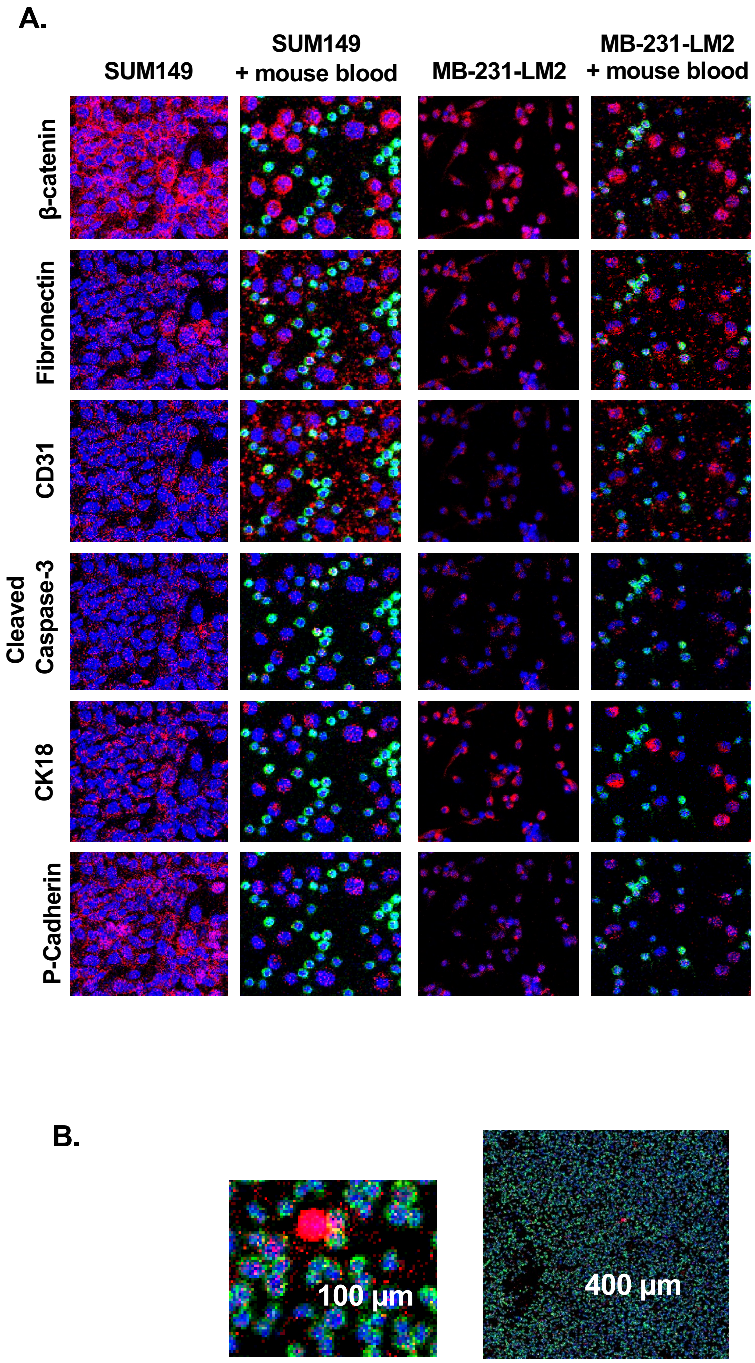


**Supplementary Figure S1: Comparison of IMC signals by specific antibodies in SUM149 and MB-231-LM2 cells with and without admixed mouse blood.** (A) Representative images showing each target in red pseudo color, DNA in blue and CD45 in green. (B) Representative examples of images at low and high magnification with scale bars.

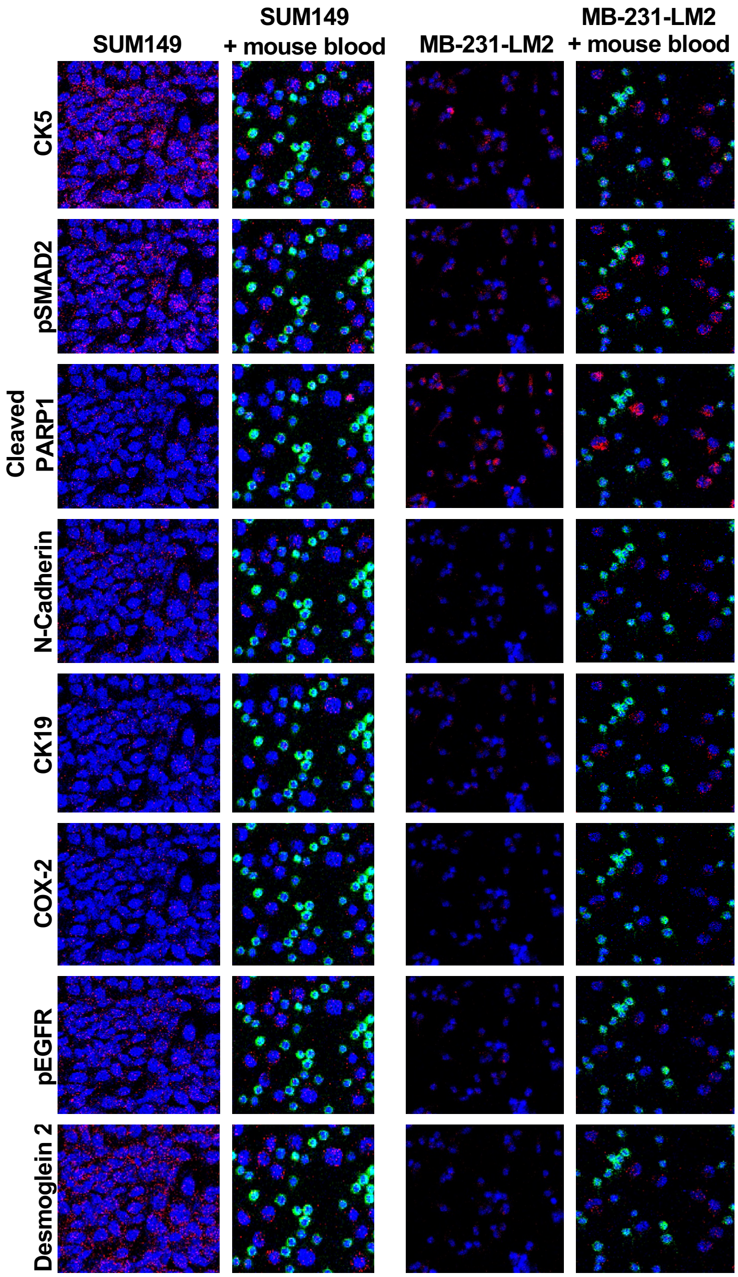


**Supplementary Figure S2: Comparison of IMC signals by specific antibodies in SUM149 and MB-231-LM2 cells with and without admixed mouse blood.** Representative images showing each target in red pseudo color, DNA in blue and CD45 in green.

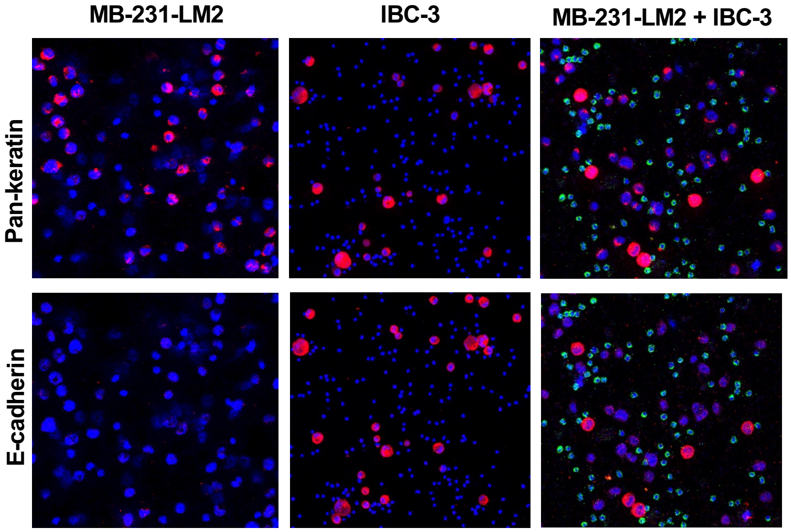


**Supplementary Figure S3: Validation of E-cadherin antibody.** Expression of PanCK and E-cadherin as detected by clone 24E10 in MDA-MB-231-LM2 and IBC-3 cells spiked into mouse blood individually (CD45 signal was omitted) or together (CD45+ staining in green).

A.

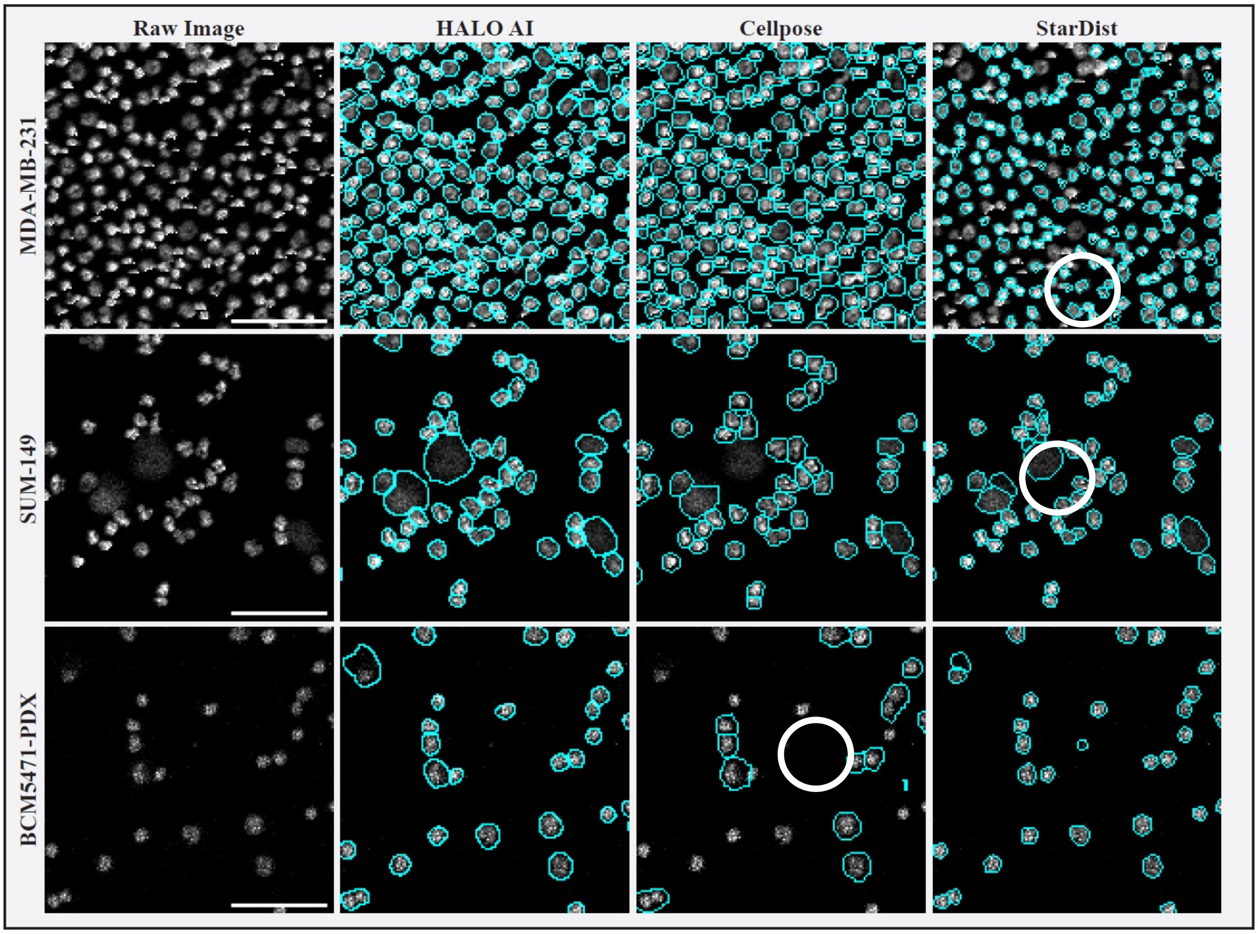


B.

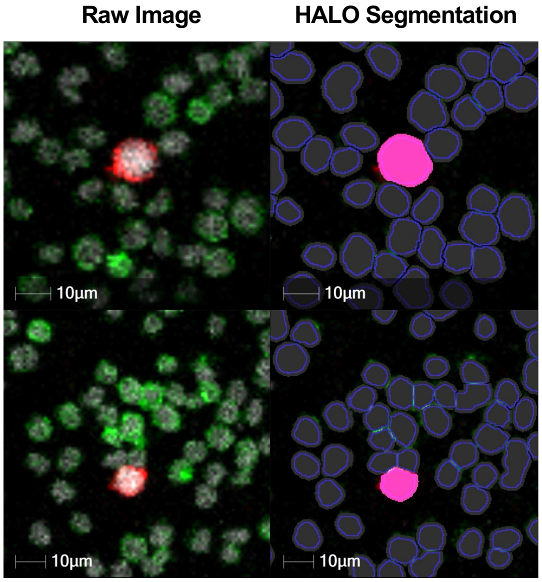


**Supplementary Figure S4. Identification of CTCs by an automated and custom AI Halo cell segmentation classifiers. A)** Nuclear segmentation of liquid biopsies from three tumor cell models across three segmentation algorithms, HALO AI, Cellpose, and StarDist (scale bars, 50µm). White circles indicate regions with errors. **B)** Example of HALO segmentation images derived from the raw data as indicated (DNA: white, PanKeratin: red, CD45: green).

### Table S1: Antibodies used or tested for IMC in this study

| Target * | Category | Metal Tag | Clone | Vendor | Catalog Number | Concentration | Comment |
|---|---|---|---|---|---|---|---|
| β-catenin | Signaling | 151Eu | 196618 | Biotechne | MAB13291-100 | 1 µg/ml | Non-specific (?) signal in blood |
| Cleaved Caspase 3 | Cell death | 172Yb | Asp175 | Standard BioTools | 3172027D | 1 µg/ml | Non-specific signal |
| CD31/PECAM-1 | Endothelial | 154Sm | 390 | Thermo Fisher | 14-0311-85 | 1 µg/ml | High background |
| CD44 | Signaling | 173Yb | 156-3C11 | CST | 3570 | 0.5 µg/ml | |
| CD45 | Mouse-specific | 175Lu | 30-F11 | Standard BioTools | 3175010B | 1 µg/ml | |
| CEBPD | Signaling | 149Sm | EPR23518-259 | Abcam | ab270410 | 1 µg/ml | No signal |
| COX2 | Signaling | 160Gd | D5H5 | CST | 73315SF | 1µg/ml | No signal |
| Cytokeratin 18 | Luminal | 155Gd | 810811 | Biotechne | MAB7619 | 0.5 µg/ml | |
| Cytokeratin 19 | Luminal | 159Tb | BA17 | Biotechne | MAB3506 | 0.5 µg/ml | |
| Cytokeratin 5 | Basal | 163Dy | monoclonal | LS Bio | LS-C812437-100 | 1 µg/ml | |
| Cytokeratin 8 | Luminal | 152Sm | LP3K | Biotechne | MAB3165 | 0.5 µg/ml | |
| Desmoglein 2 | Epithelial | 146Nd | 6D8 | Biotechne | LS-C761566-100 | 1 µg/ml | No signal |
| E-cadherin | Epithelial | 148Nd | 180224 | Biotechne/CST | MAB18381/3195 | 0.5 µg/ml | No signal |
| E-cadherin | Epithelial | 158Gd | 24 E10 | Standard BioTools | 3158029D | 1 µg/ml | |
| EGFR | Signaling | 144Nd | AY13 | Biolegend | 352902 | 1 µg/ml | |
| EGFR (pY1068) | Signaling | 162Dy | D7A5 | CST | 48576SF | 1 µg/ml | No signal |
| EpCAM | Epithelial | 147Sm | 9C4 | Biolegend | 324202 | 1 µg/ml | |
| Fibronectin | Mesenchymal | 153Eu | EPR23110-46 | Abcam | ab268022 | 1 µg/ml | High background |
| Lamin B1 | Human-specific | 12G6 | HS-404 017 | SYSY Antibodies | HS-404 017 | 1 µg/ml | |
| MHC Class II | Human-specific | 154Sm | TDR31.1 | LS Bio | LS-B6315-50 | 1 µg/ml | |
| N-cadherin | Mesenchymal | 158Gd | CDH2/1573 | LS Bio | LS-C761774-100 | 1 µg/ml | |
| Na/K ATPase | Human-specific | 170Er | D4Y7E | CST | 23565S | 1 µg/ml | No signal |
| NOS2 | Signaling | 145Nd | 4 E5 | Novus | NBP2-22119 | 1 µg/ml | |
| NUMA1 | Human-specific | 141Pr | SPM300 | LS Bio | LS-C390797-100 | 1 µg/ml | High background |
| PARP1, cleaved | Cell death | 168Er | E51 | Abcam | ab203467 | 1 µg/ml | Non-specific signal in MB231 |
| P-cadherin | Epithelial | 161Dy | 106020 | Biotechne | MAB761-100 | 1 µg/ml | Non-specific/low signal |
| Pan-keratin | E/M | 169Tm | Polyclonal | Thermo Fisher | 26411-1-AP | 0.33 µg/ml | |
| SMAD2 | Signaling | 167 Er | EP784Y | Abcam | ab157371 | 1 µg/ml | |
| SMAD2 (pS467) | Signaling | 165Ho | EPR23681-40 | Abcam | ab280897 | 1 µg/ml | Low signal |
| Vimentin | Mesenchymal | 150Nd | RV203 | Abcam | ab8979 | 0.5 µg/ml | |
| ZEB1 | Mesenchymal | 164Dy | EPR17375 | Abcam | ab228986 | 1 µg/ml | |

*Green shade indicates antibodies that were considered working well based on integration of results across cell line models and staged approaches.

**Table S2: Comparison of the frequency of CTCs in blood collected by tail vein versus cardiac puncture.**

| Mouse I.D. | Cardiac | Tail vein | TV ($mm^3$) |
|---|---|---|---|
| SUM149 #1 | 12 | n.d. | 3704.8 |
| SUM149 #2 | 2 | n.d. | 3269.3 |
| SUM149 #3 | 0 | n.d. | 2572.5 |
| SUM149 #4 | 4 | 1 | 2178.6 |
| SUM149 #5 | 0 | 1 | 2384.8 |
| SUM149 #6 | 1 | 0 | 2151.7 |
| MB-231-LM2 #1 | 0 | n.d. | 2100 |
| MB-231-LM2 #2 | 12 | n.d. | 2427.6 |
| MB-231-LM2 #3 | 6 | n.d. | 1825.2 |
| MB-231-LM2 #4 | 0 | 1 | 2112.0 |
| MB-231-LM2 #5 | 6 | 2 | 3447.8 |
| MB-231-LM2 #6 | 15 | 4 | 2788.2 |
| BCM5471 #1 | 3 | n.d. | 2048.0 |
| BCM5471 #2 | 0 | n.d. | 2427.6 |
| BCM5471 #3 | n.d. | 3 | 2573.5 |
| BCM5471 #4 | 37 | 2 | 1825.9 |
| BCM5471 #5 | 10 | 16 | 2573.5 |
| BCM5471 #6 | 18 | 6 | 2664.4 |

Manual count of the number of CTCs per 100 uL of blood collected from 6 mice each per tumor model (n.d., not done; TV, tumor volume).

**Table S3: Threshold parameters for calling PanKeratin+ CTCs by HALO HighPlex FL module per tumor model and instrument.**

| Tumor model | Instrument | DNA3 | PanKeratin* | CD45* |
|---|---|---|---|---|
| BCM5471 | Hyperion+ | ≥ 7 | ≥ 1.5 (40%) | < 1.5 (25%) |
| BCM5471 | Hyperion XTI | ≥ 3 | ≥ 2.75 (35%) | < 1.5 (25%) |
| MB-231-LM2 | Hyperion+ | ≥ 10 | ≥ 3 (45%) | < 1.5 (25%) |
| MB-231-LM2 | Hyperion XTI | ≥ 1.75 | ≥ 3 (50%) | < 1.5 (25%) |
| SUM149 | Hyperion+ | ≥ 7 | ≥ 1.8 (40%) | < 1.5 (25%) |
| SUM149 | Hyperion XTI | ≥ 10 | ≥ 6 (50%) | < 1.5 (25%) |

*Signal intensity thresholds and % completeness, i.e. percentage of the cell area covered by signal.

**Table S4: Comparison of CTC counts by HALO compared to manual counts.**

| Mouse Model Sample# | Manual 1 | Manual 2 | HALO |
|---|---|---|---|
| SUM-149 #1 | 15 | 12 | 14 |
| SUM-149 #2 | 2 | 2 | 5 |
| SUM-149 #3 | 0 | 0 | 1 |
| SUM-149 #4 | 1 | 1 | 0 |
| SUM-149 #5 | 3 | 4 | 3 |
| SUM-149 #6 | 0 | 0 | 0 |
| SUM-149 #7 | 4 | 1 | 2 |
| SUM-149 #8 | 0 | 0 | 1 |
| SUM-149 #9 | 2 | 1 | 1 |
| MB-231-LM2 #1 | 0 | 0 | 1 |
| MB-231-LM2 #2 | 0 | 12 | 1 |
| MB-231-LM2 #3 | 4 | 6 | 3 |
| MB-231-LM2 #4 | 2 | 2 | 2 |
| MB-231-LM2 #5 | 16 | 15 | 14 |
| MB-231-LM2 #6 | 1 | 4 | 0 |
| MB-231-LM2 #7 | 0 | 0 | 1 |
| MB-231-LM2 #8 | 9 | 9 | 8 |
| MB-231-LM2 #9 | 6 | 6 | 4 |
| MB-231-LM2 #10 | 3 | 3 | 3 |
| BCM-5471 #1 | 3 | 3 | 4 |
| BCM-5471 #2 | 0 | 3 | 3 |
| BCM-5471 #3 | 0 | 0 | 0 |
| BCM-5471 #4 | 23 | 37 | 26 |
| BCM-5471 #5 | 0 | 2 | 0 |
| BCM-5471 #6 | 2 | 3 | 3 |
| BCM-5471 #7 | 6 | 10 | 9 |
| BCM-5471 #8 | 3 | 16 | 8 |
| BCM-5471 #9 | 17 | 18 | 18 |
| BCM-5471 #10 | 0 | 6 | 2 |

CTC frequency (#CTC/100uL blood) as assessed in the same ROI by two independent investigators and as determined by HALO.
